# Inhibition of mitochondrial ferredoxin 1 (FDX1) prevents adaptation to proteotoxic stress

**DOI:** 10.1101/288365

**Authors:** Peter Tsvetkov, Alexandre Detappe, Kai Cai, Heather R. Keys, Zarina Brune, Weiwen Ying, Prathapan Thiru, Mairead Reidy, Guillaume Kugener, Aviad Tsherniak, Sandro Santagata, Luke Whitesell, John L. Markley, Irene M. Ghobrial, Susan Lindquist

**Affiliations:** Whitehead Institute for Biomedical Research, Cambridge, MA 02142; Dana-Farber Cancer Institute, Harvard Medical School, Boston, MA 02215; Biochemistry Department, University of Wisconsin-Madison, Madison, WI 53706; OnTarget Pharmaceutical Consulting LLC, Lexington MA 02421; Broad Institute of Harvard and MIT, Cambridge, MA 02142; Department of Pathology, Brigham and Women’s Hospital, Harvard Medical School, Boston, MA 02115; Howard Hughes Medical Institute, Department of Biology, Massachusetts Institute of Technology, Cambridge, MA 02139

**Author notes:** Deceased (Susan Lindquist).

## Abstract

The mechanisms used by cancer cells to resist the severe disruption in protein homeostasis caused by proteasome inhibitors remain obscure. Here, we show this resistance correlates with a metabolic shift from glycolysis to oxidative phosphorylation (OXPHOS). Employing small molecule screens, we identified a striking overlap between compounds that preferentially impede the growth of proteasome inhibitor-resistant cancer cells and those that block the growth of high OXPHOS cells. Elesclomol potently exhibits both characteristics. Using genome-wide CRISPR/Cas9-based screening, in vitro validation and NMR spectroscopy we identify mitochondrial protein ferredoxin 1 (FDX1), a critical component of mitochondrial iron-sulfur (Fe-S) cluster biosynthesis, as the primary target of elesclomol. In a mouse model of multiple myeloma, inhibition of FDX1 with elesclomol significantly attenuated the emergence of proteasome inhibitor-resistance and markedly prolonged survival. Our work reveals that the mitochondrial Fe-S cluster pathway is a targetable vulnerability in cancers that are resistant to increased proteotoxic burden.

## INTRODUCTION

Cancer cells develop resistance to the cytotoxic effects of chemotherapy by multiple mechanisms, including acquired mutations, gene amplification, re-wiring of transcriptional networks, and shifting to an altered cell state (Easwaran et al., 2014; Holohan et al., 2013). Much effort to understand the emergence of drug resistance has focused on the identification of acquired mutations of the proximal therapeutic target itself as exemplified in the setting of kinase inhibitor-resistance (Gorre et al., 2001; Paez et al., 2004). However, alternative mechanisms of drug resistance may arise independently of genetic mutations. In such instances, there may be selection of a pre-existing population of cells that has an intrinsically resistant state or the treatment may induce a cell-state switch that endows cancer cells with the ability to withstand the insult and persist (Meacham and Morrison, 2013; Schmitt et al., 2016). The initial ability to persist following treatment may also involve a combination of these mechanisms, with a drug resistant cell-state fostering conditions required for the accumulation of additional mutations that confer a stable state of unfettered drug-resistance (Hata et al., 2016; Ramirez et al., 2016). Thus, an immediate critical need exists for diverse strategies to limit the emergence of distinct drug resistance. In addition to the immediate importance of developing strategies to limit the emergence of specific drug resistant states, understanding the mechanisms through which cancers withstand the initial toxic effects of treatment can reveal specific targetable oncogenic dependencies.

Recent studies show that drug-resistance in several cancer types is associated with altered cell metabolism (Farge et al., 2017; Ippolito et al., 2016; Kuntz et al., 2017; Lee et al., 2017; Matassa et al., 2016; Vazquez et al., 2013). Cancers are known to rewire their metabolic networks to support the energetic and biosynthetic demands for proliferation (Cairns et al., 2011; Vander Heiden et al., 2009). While this is typically associated with aerobic glycolysis (Warburg effect), mitochondrial respiration is also required for tumor proliferation (Vyas et al., 2016; Weinberg et al., 2010). This mitochondrial dependency involves functions beyond ATP production, including generation of reactive oxygen species (ROS), maintenance of cellular redox balance, amino acid biosynthesis and more (Cantor et al., 2017; Cantor and Sabatini, 2012; Sullivan et al., 2015; Zong et al., 2016). Yet, how such metabolic reprograming contributes to drug resistance is still largely unknown.

Proteasome inhibition is a well-established example of an initially efficacious therapeutic strategy that is rendered ineffective by intrinsic and acquired drug-resistance. Proteasome-mediated protein degradation is a key regulator of protein homeostasis (Collins and Goldberg, 2017; Deshaies, 2014; Labbadia and Morimoto, 2015; Tanaka et al., 2012) and the increased flux of proteins in cancer results in a greater dependence on proteasome function (Kumar et al., 2012; Luo et al., 2009; Petrocca et al., 2013). This dependency has been exploited with proteasome inhibitors to specifically target cancer cells in experimental models and in the clinic as a frontline therapy for multiple myeloma.

The therapeutically-employed proteasome inhibitors target the catalytic activity of the proteasome. Paradoxically, cancer cells in culture are extremely sensitivity to proteasome inhibitors, yet most tumors in patients readily withstand the proteotoxic effects of these inhibitors. Thus, the clinical utility of proteasome inhibitors is greatly restricted (Manasanch and Orlowski, 2017). There were several suggested mechanisms of resistance including constitutive activation of NF-κB (Markovina et al., 2008) and the chaperone machinery (Li et al., 2015) and alterations in the EGFR/JAK1/STAT3 pathway (Zhang et al., 2016). Moreover, a predominant hypothesis was that cancer cells develop resistance to proteasome inhibitors by acquiring mutations in the catalytic sites targeted by the proteasome inhibitors (Kisselev et al., 2012). Although such mutations have been detected in tissue culture-derived resistance models (Oerlemans et al., 2008; Ri et al., 2010), they are absent in clinical samples (Lichter et al., 2012). Therefore, alternative resistance mechanisms must exist.

Several recent studies using different genetic approaches have converged on a novel mechanism that allows cells to cope with the proteotoxic stress induced by proteasome inhibitors (Acosta-Alvear et al., 2015; Shi et al., 2017; Tsvetkov et al., 2015; Tsvetkov et al., 2017). The 26S proteasome complex consists of the catalytic barrel where proteins are processed (20S complex) that can be capped with either one or two 19S regulatory complex caps that include the ubiquitin-regulating enzymes and protein unfolding ATPases (Budenholzer et al., 2017; Coux et al., 1996). Suppressing the expression of any one of the many subunits of this 19S complex results in a proteasome inhibitor-resistant state, which we refer to as the Low 19S Subunit (Lo19S) state. When the expression of a 19S subunit is suppressed, levels of the intact 26S proteasome complex are decreased, resulting in a greater proportion of 20S complexes. However, the overall cellular capacity for proteasome-mediated degradation is unimpaired (Tsvetkov et al., 2015). In this Lo19S state, cells exhibit an enhanced ability to tolerate the toxic effects of many proteasome inhibitors. This resistance mechanism has been observed in experimental model systems and in a variety of tumors recovered from patients (Acosta-Alvear et al., 2015; Nijhawan et al., 2012; Tsvetkov et al., 2017). Prominently, multiple myeloma patients that are refractory to proteasome inhibitors most often have tumors with the spontaneously reduced expression of 19S subunits (Acosta-Alvear et al., 2015; Tsvetkov et al., 2017). These findings suggest that the Lo19S state is a frequent, naturally occurring mechanism that cancer cells deploy to resist the toxic effects of proteasome inhibition.

In this study, we use a functional genomics approach to demonstrate that the proteasome inhibitor resistant, Lo19S state is coupled to a metabolic shift that increases dependence on OXPHOS in many tumor types revealing an actionable metabolic vulnerability. We further demonstrate that the first-in-class Ferredoxin 1-specific inhibitor, elesclomol, suppresses the shift from glycolysis to OXPHOS. Importantly, inhibition of FDX1 attenuates the ability of cancer cells to cope with proteasome inhibitor-induced toxicity both in culture and in an orthotopic mouse model of multiple myeloma.

## RESULTS

### The Lo19S state is associated with increased OXPHOS in many tumor types

To identify cellular adaptations that are associated with the Lo19S state in resected human cancer specimens, we searched for samples within The Cancer Genome Atlas (TCGA) that exhibited this state. The Lo19S state is defined by having the expression of at least one 19S proteasome subunit gene suppressed by more than 3 standard deviations from the mean (Tsvetkov et al., 2017). Parsing the genes that were differentially expressed within the Lo19S group as compared to control (Figure 1A), we found that, within breast cancer samples, 2.8% (n=31) of the tumors resided in the Lo19S state. Using gene ontology (GO) and gene set enrichment analysis (GSEA) methods to compare differential gene expression between Lo19S state breast tumors and the remaining breast tumors in the dataset, we identified a striking enrichment of networks related to mitochondrial OXPHOS processes (Figure 1B-D). We further assessed additional TCGA dataset tumors of diverse tissue origins for which sufficient numbers of Lo19S samples (>20) existed. With the single exception of low-grade glioma, all other Lo19S tumor types, including skin, thyroid, kidney and prostate cancers, were highly enriched for mitochondria-related gene categories (Figure S1A). Thus, the Lo19S state is associated with increased expression of genes associated with mitochondrial function in a variety of tumor types. The elevated OXPHOS gene signature is hypothesized to result from high OXPHOS metabolism and increased mitochondrial dependence in these tumors.

**Figure 1.**
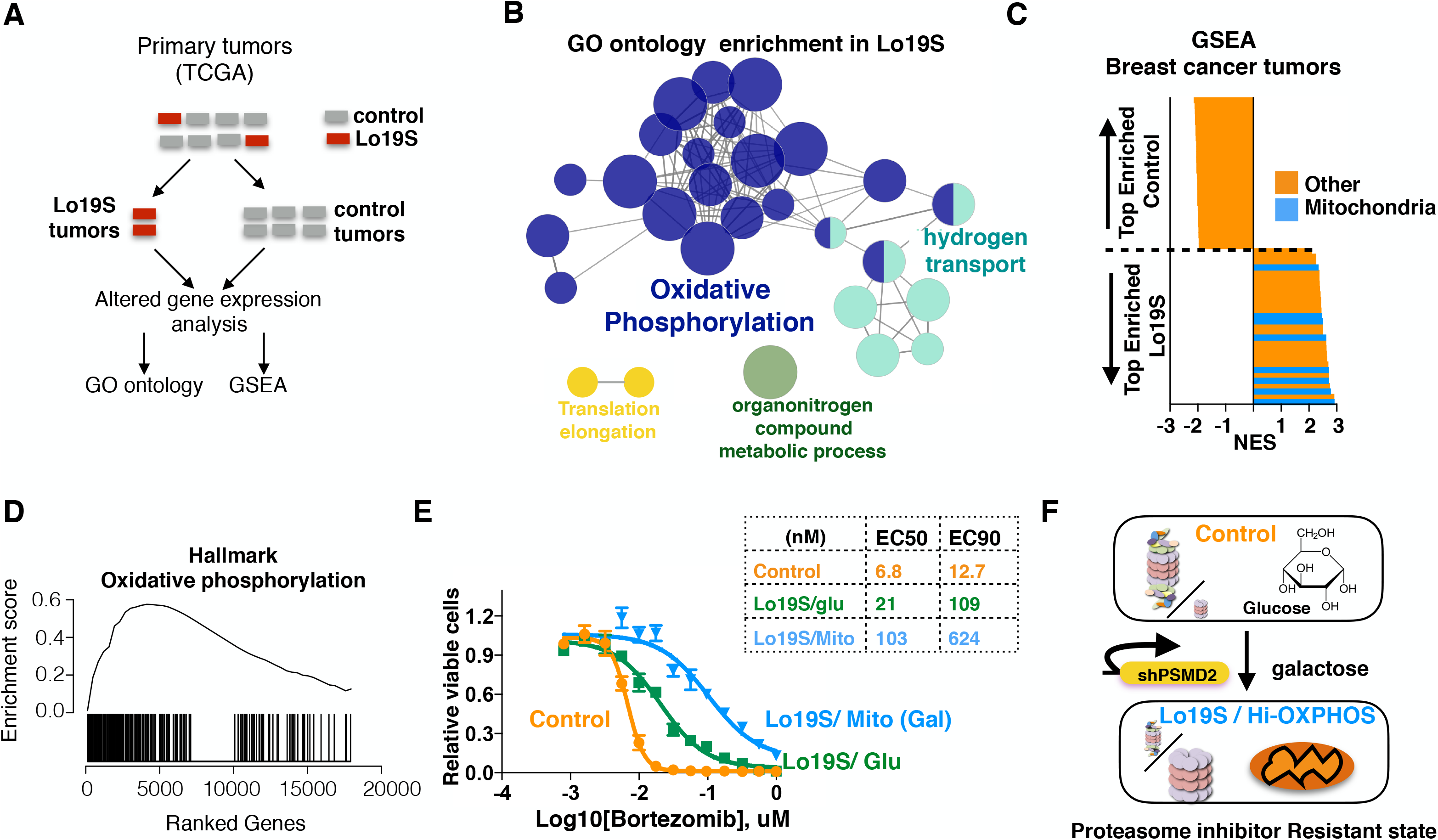
Mitochondrial respiration is associated with increased resistance to proteasome inhibitors in the Lo19S state. (A) Gene expression in cancers in the TCGA dataset was analyzed and the data stratified by tumors exhibiting the Lo19S state (one subunit of the 19S proteasome complex suppressed by more than 3 standard deviations (Tsvetkov et al., 2017)) or control (the rest of the tumors). This enables exploration of the unique gene expression signature of the Lo19S state. (B) GO ontology gene networks most enriched in the Lo19S state in breast cancer tumors (BRCA-TCGA) are plotted and color coded by function. (Log2 score >0.5, plotted using cyttoscape clue go) (C) Gene set enrichment analysis (GSEA) of genes upregulated in Lo19S but not control breast cancer tumors derived from the TCGA. Top and bottom 29 categories are plotted. Mitochondrial-associated categories are marked in blue, the remainder in orange. (D) Specific GSEA scores of genes from the mitochondrial-related category from the analysis described in (C) are ranked (hallmark of oxidative phosphorylation) (E) T47D breast cancer cells harboring a doxycycline-inducible PSMD2 shRNA were grown in the presence or absence of 0.2 μg/ml doxycycline for 72 hours to induce the Lo19S state. Cells were then collected, washed and plated in the absence of doxycycline in media containing either glucose (Glu) or galactose. Galactose induces mitochondrial respiration (Mito). The relative viability was measured 72 hours after addition of the indicated concentrations of bortezomib. The calculated EC50s and EC90s are also presented. (F) Schematic showing the shift to the Lo19S state (shPSMD2) and mitochondrial respiration (galactose instead of glucose) representing the proteasome inhibitor resistant state.

To directly test whether altered cellular metabolism can modulate the sensitivity of cancer cells to proteasome inhibitors, we engineered the breast cancer cell line T47D into Lo19S and increased OXPHOS states. Initially, we reconstituted the Lo19S state by transient suppression of the PSMD2 subunit of the 19S complex using an inducible shRNA. As expected, transiently inducing the Lo19S state resulted in increased resistance to the chemically distinct proteasome inhibitors bortezomib (Figure S1B) and Ixazomib (Figure S1C) increasing their proliferation inhibition EC50s by almost 4-fold and their EC90s by 14-27-fold (Figure S1B and S1C).

To induce a high OXPHOS (Hi-OXPHOS) state, we changed the carbon source in the cell culture media from glucose to galactose, which is poorly fermentable. This method has been previously shown to drive glutamine utilization by mitochondria and to thus increase mitochondrial respiration (Gohil et al., 2010). Interestingly, enhancing respiration alone caused mild resistance to proteasome inhibition (Figure S1D), as well as reduced heat-shock induction by proteasome inhibitors (Figure S1E) without affecting the overall catalytic activity of cellular proteasomes (Figure S1F). While these results support an association between the cellular metabolic state and the response to proteasome inhibition, the effects of inducing a Hi-OXPHOS state were comparatively mild to the resistance achieved by inducing the Lo19S state. We reasoned that the Hi-OXPHOS signature we identified in Lo19S patient tumors may cooperate with the Lo19S state to further promote the ability of the cells to withstand proteotoxic stress. Indeed, increasing respiration in cells in the Lo19S state further increased the Lo19S-induced resistance to proteasome inhibitor-induced cell death by approximately an additional 5-10-fold (Figure 1E and S1G). Strikingly, combining the Lo19S and Hi-OXPHOS states results in a 50-fold increase in the EC90 for Bortezomib (Figure 1E and S1G). Thus, induction of the Hi-OXPHOS state increases the ability of cells to withstand proteasome inhibitor induced proteotoxic stress, particularly in the background of the Lo19S state. (Figure 1F).

### Elesclomol preferentially targets the proteasome inhibitor resistant and Hi-OXPHOS states

A deeper understanding of the proteasome inhibitor resistant state could yield insights into the molecular mechanisms activated by cells to cope with proteotoxic stress and elucidate new therapeutic strategies for cancer. With these goals in mind, we screened mechanistically diverse small molecule libraries to identify compound sensitivities altered by the Lo19S and Hi-OXPHOS states. An initial screen was conducted in the isogenic T47D model with transient induction of the Lo19S state as described above. We evaluated >4300 mechanistically well-defined compounds within 4 libraries. The compounds included approved cancer drugs (Selleck anti-cancer library: **CDL**), common natural products (Natural product library: **NPC**) and an NIH bioactives library (**NIH)**. Each of these compounds was tested at ≥ 4 concentrations. (**Table S1**). An additional 2866 compounds from the Boston University’s Chemical Methodology and Library Development (CMLD-BU) small molecule collection was screened at a single dose (**BUCL**, see **Table S1**). Across the four independently performed small molecule screens, the Lo19S state conferred resistance almost exclusively to proteasome inhibitors (Figure 2A, S2A, **Table S2**). Interestingly, the Lo19S state selectively sensitized cells to only a very small number of compounds. These included elesclomol, disulfiram and the pro-apoptotic agent ABT-263 (Figure 2A, **Table S2**). The small number of hits suggests that in our cellular system the inducible Lo19S state does not induce a global rewiring of drug sensitivities but rather exerts a very specific effect that results in resistance to multiple distinct proteasome inhibitors and sensitivity to only a hand full of compounds.

**Figure 2.**
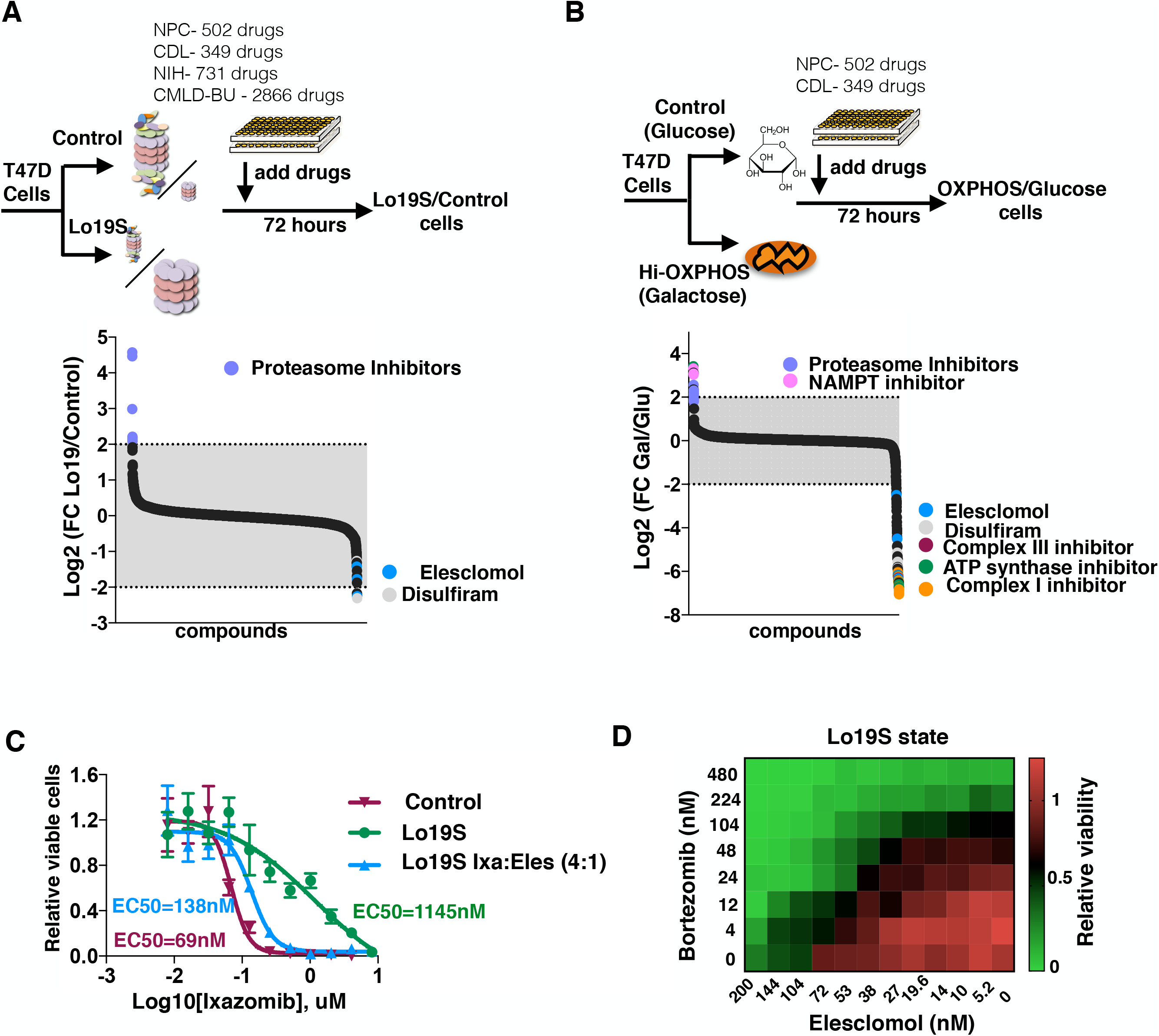
Elesclomol targets the Lo19S state to block proteasome inhibitor resistance. (A) T47D inducible Lo19S state cells or control cells (as described in Figure 1E) were exposed to 349 compounds from the Selleck anti-cancer L3000 drug library (CDL) at 4 concentrations, the natural product library (NPC) comprising 502 compounds at 5 concentrations and the NIH bioactive compound library that includes 731 compounds in 4 concentrations. The relative viability of the Lo19S versus control cells was calculated for replica experiments and the log2 of the ratio is plotted. (B) T47D cells were grown in the presence of either glucose or galactose as the carbon source in the presence of 349 compounds from the Selleck anti-cancer L3000 drug library (CDL) in 4 doses and the natural product library (NPC) comprising 502 compounds in 5 doses. The relative viability of the respiring cells (Hi-OXPHOS) (gal) versus the control cells (glu) was calculated for replica experiments and the log2 of the ratio is plotted. Specific compounds are annotated. (C-D) Elesclomol re-sensitizes Lo19S T47D cells to proteasome inhibition. C) The effect on relative cell growth of elesclomol added with the proteasome inhibitor ixazomib at a 1:4 (elesclomol:ixazomib) ratio (the ratio of the EC50s) to Lo19S cells compared to the effect of ixazomib alone added to either Lo19S or control cells. Plotted are the mean +-SD of at least three replicas and the calculated EC50s. D) Heat map of relative cell growth following addition of different combinations of bortezomib and elesclomol to the Lo19S state cells.

We conducted a second set of drug screens to identify unique sensitivities conferred by the Hi-OXPHOS state using parental T47D breast cancer cells (without 19S subunit suppression). For this screen, we used the NPC and CDL libraries as they contained the greatest number of hits in the Lo19S screen. We tested over 800 compounds at multiple concentrations and measured their effects on relative cell growth/viability when cells were grown under aerobic glycolysis conditions (glucose-supplemented medium) or forced mitochondrial respiration conditions (galactose-supplemented medium). Interestingly, several different proteasome inhibitors (bortezomib, MLN9708, MLN2238) and the nicotinamide phosphoribosyltransferase (NAMPT) inhibitor, APO866, were the strongest hits in this screen, scoring highly across multiple doses (Figure 2B). In contrast, when the same panel of compounds was tested in cells grown in galactose (Hi-OXPHOS state), perhaps unsurprisingly, we found increased sensitivity to compounds that target mitochondrial respiration including Complex I (Rotenone, Degeulin and QNZ), Complex III (antimycin A), and ATP synthase (Oligomycin A) inhibitors. Surprisingly, however, the small set of compounds that preferentially targeted respiring cells across multiple doses and that are not known to directly target the mitochondria respiratory chain included both elesclomol (5 concentrations) and disulfiram (4) (Figure 2B and **Table S2**).

Strikingly, the results of our two different screening strategies virtually mirrored one another: proteasome inhibitors were more potent in glycolytic cells compared to Hi-OXPHOS cells and less effective in the context of the Lo19S state. Conversely, elesclomol and disulfiram were more potent in cells in the Hi-OXPHOS state compared with glycolytic cells and were highly effective in slowing the growth of cells in the Lo19S state. The drug sensitivity and resistance profiles of these cell states suggest that induction of either the Lo19S state or the Hi-OXPHOS state affects the cell similarly but not identically.

The fact that other respiratory chain inhibitors were not identified in the Lo19S drug screens suggests that elesclomol and disulfiram work by a distinct mechanism of action compared to other known respiratory chain inhibitors. The ability of a compound to preferentially target the Lo19S state might be extraordinarily beneficial in the context of proteasome inhibitor resistance, we tested whether elesclomol could re-sensitize the Lo19S cells to proteasome inhibitors. When we exposed cancer cells to the combination of elesclomol and proteasome inhibitors, we observed a synergistic interaction that eliminated the relative resistance of cells in the Lo19S state to proteasome inhibitors (Figure 2C and 2D).

Consistent with prior studies on the anti-cancer activity of elesclomol (O’Day et al., 2009), we found that it is a potent inhibitor of cancer cell growth across a variety of tumor types (Figure 3A). In addition to the intrinsic sensitivity of certain cancer cells to elesclomol, as our drug screen indicated, the sensitivity of cancer cells to elesclomol can be further enhanced by forcing them into a Hi-OXPHOS state. This phenomenon is observed using mitochondrial targeting agents such as antimycin A (Figure S3A). In some cases, shifting to OXPHOS conditions can also lower the EC50 of elesclomol by two to three orders of magnitude (Figure 3B).

**Figure 3.**
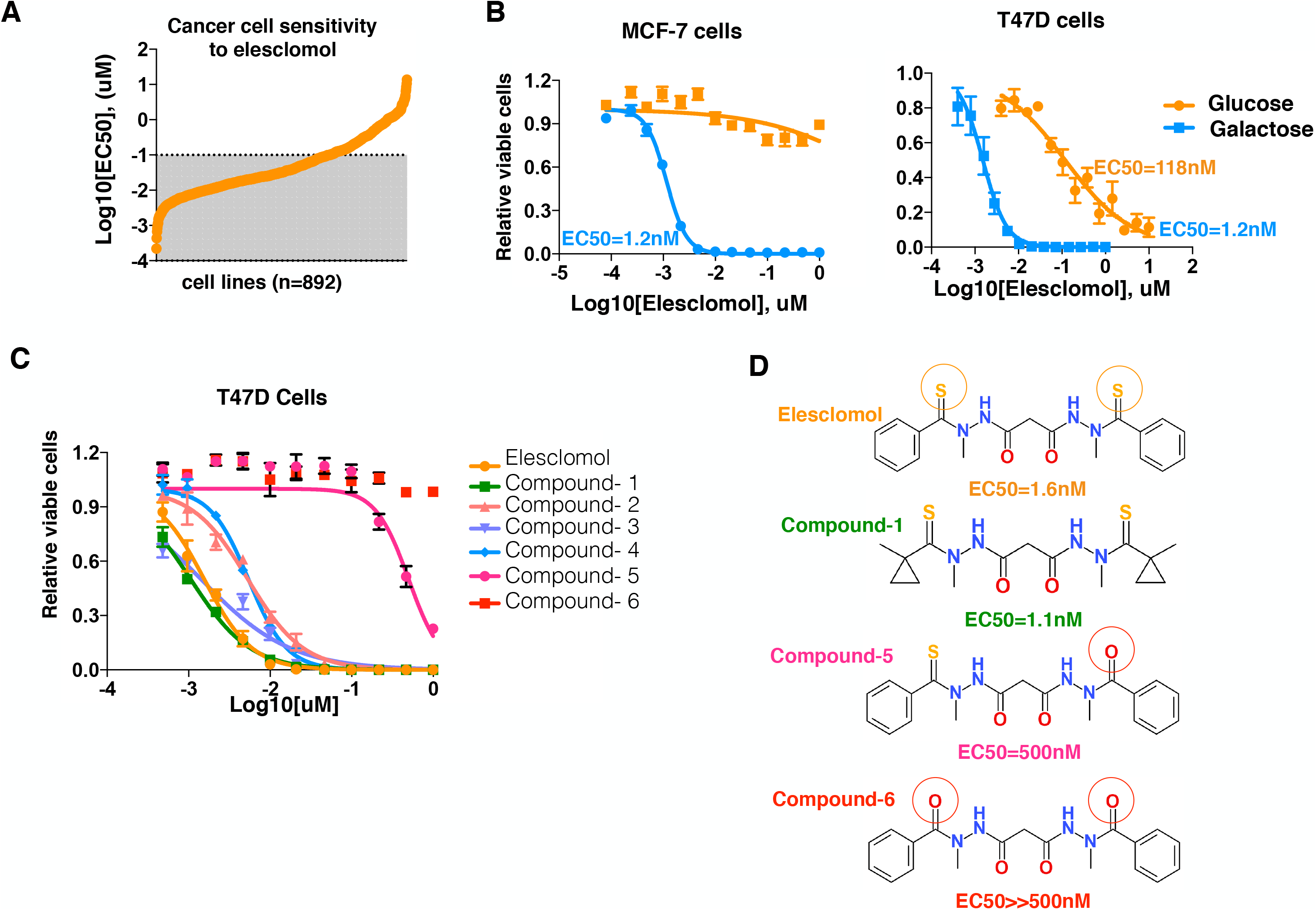
Elesclomol preferentially targets cells in the Hi-OXPHOS state. (A) The EC50 of elesclomol for 892 cancer cell lines as taken from the GDSC dataset (http://www.cancerrxgene.org/). (B) MCF7 and T47D cells were grown in the presence of either glucose (control, orange) or galactose (Hi-OXPHOS, blue) as the carbon source and the relative cell number was analyzed 72 hours after addition of the indicated concentrations of elesclomol. (C) T47D cells were grown in the presence of galactose (Hi-OXPHOS) and exposed to various elesclomol analogs (compounds 1-6).The relative cell growth (color coded) was analyzed 72 hours post compound addition. Plotted are the mean +-SD of at least three replicas and the calculated EC50s (B-C) (D) Specific analogs of elesclomol. Reactive groups are indicated with a colored circle.

We examined several elesclomol analogs (Figure S3B) to determine the structural features that are important for mediating its cytotoxicity. For example, elesclomol possesses two sulfur groups that presumably play a role in its bioactivity. Substitution of oxygen for even one of the sulfur groups (compound-5) reduced the activity of elesclomol by 300 fold. The loss of activity was even more pronounced (>> 300 fold) when both sulfurs were substituted (compound-6) (Figure 3C, 3D and S3C). Various other modifications to the elesclomol scaffold were tolerated, as long as the core sulfur geometry remained intact.

### Ferredoxin 1 (FDX1), a component of the iron-sulfur cluster assembly pathway, is the primary mediator of elesclomol-induced toxicity

Despite its use in clinical trials on a limited number of malignancies, the elesclomol mechanism of action remains unclear. We therefore undertook a genetic approach to identify its direct target and to define cellular pathways that modulate the sensitivity of cancer cells to elesclomol-induced toxicity. Two elesclomol analogs that are potent inhibitors of K562 cell growth (compounds **1** and **2**) were chosen for this genetic screen (Figure S4A-B). We performed a genome-wide, positive selection CRISPR/Cas9-based screen to identify genes whose loss conferred resistance to these elesclomol analogs (Figure 4A) using an activity-optimized library consisting of 187,535 sgRNAs targeting 18,633 protein-coding genes (Wang et al., 2015). We transduced K562 cells with the sgRNA library (Wang et al., 2016), and passaged the pool of CRISPR/Cas9-targeted cells in the presence of different concentrations of compound-**1** or -**2** over 30 days to maintain the cells in an actively proliferating state. We measured the relative abundance of each sgRNA in the compound-treated K562 cells at the beginning and end of the culture period. For each gene, we calculated its score as the mean log_2_ fold-change in abundance of all sgRNAs targeting the gene. In both screens, a single gene target scored significantly higher than any other gene: *FERREDOXIN 1*, (*FDX1)* (Figure 4B and 4C). To validate FDX1 as a target of elesclomol, we created an *FDX1*-null cell line model using two individual sgRNAs or a non-targeting control sgRNA. Indeed, loss of FDX1 protein (Figure 4D) confers relative resistance to elesclomol (Figure 4E). If FDX1 is a key target of elesclomol, then *FDX1* deletion may limit the ability of cells to shift their core energy metabolism from aerobic glycolysis to mitochondrial respiration. We tested this hypothesis using our *FDX1* knock-out cell lines. Indeed, cells deleted for *FDX1* were not able to make the metabolic shift that enables growth in the absence of glucose (Figure 4G and 4H). Altogether, these results establish FDX1 as a critical target of elesclomol that is necessary for the glycolysis-to-OXPHOS switch.

**Figure 4.**
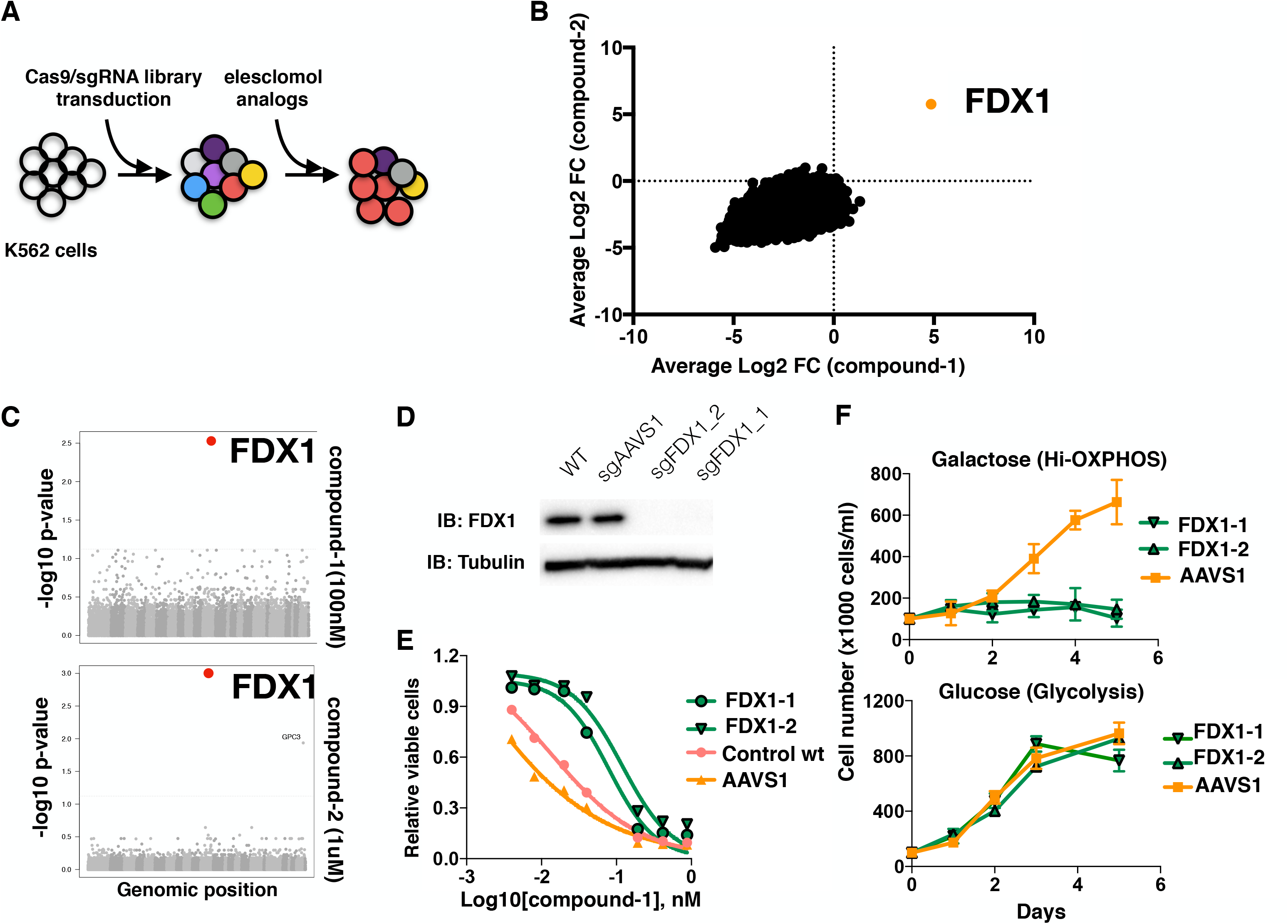
CRISPR/Cas9 genetic screen analyses reveal FDX1 as a mediator of elesclomol-induced toxicity. (A) Schematic depicting the pooled CRISPR-based screen. (B) Gene scores in elesclomol-1-(100 nM) and elesclomol-2-(1uM) treated K562 cells. The gene score is the median log2 fold change in abundance of all sgRNAs targeting that gene during the culture period. The FDX1 score is indicated. (C) The corrected p-values (-log10) of the KS tests of the sgRNA distribution for each gene vs the distribution of all sgRNAs in the screen in the eleslcomol-1 and elesclomol-2 screens. Values are ordered on the x-axis by chromosome and location; the dotted line indicates a corrected p-value of 0.05. The FDX1 score is indicated. (D) Western blot analysis of FDX1 and tubulin (loading control) protein expression levels in WT K562 cells (WT) or cells with FDX1 (two distinct sgRNAs) and AAVS1 deletions. (E-F) Viability curves of parental K562 cells and cells deleted for either AAVS1 (control) or FDX1 achieved with two sgRNAs using CRISPR/Cas9. (E) The indicated cells were treated with increasing concentrations of eleslcomol-1 and viability was examined after 72 hours. (F) The indicated cells were grown in the presence of either glucose or galactose and the relative cell number plotted.

### Elesclomol directly binds and inhibits FDX1 function to block iron-sulfur cluster formation

FDX1 is active in both iron-sulfur (Fe-S) cluster formation and the steroid hormone synthesis pathway (Cai et al., 2017b; Lill and Muhlenhoff, 2006; Sheftel et al., 2010). We examined genome-scale loss-of-function CRISPR/Cas9 screens from the Cancer Dependency Map (Meyers et al., 2017) to identify genes whose knockout phenocopies elesclomol sensitivity across hundreds of cancer cell lines. We found that cell lines that are sensitive to elesclomol show higher sensitivity to the knockout of genes involved in Fe-S cluster assembly (Figure S4C, **Table S3**).. This finding strongly supports our identification of FDX1 as a key modulator of sensitivity to elesclomol, and indicates its role in Fe-S cluster assembly as its critical function in the context of elesclomol treatment.

Fe-S clusters are generated from sulfur extracted from cysteine by the iron sulfur-cluster (ISC) core complex (NFS1-ISD11-Acp-ISCU)_2_ and iron. A reductant (FDX1 or FDX2) is required for Fe-S cluster biosynthesis (Lill and Muhlenhoff, 2006; Py and Barras, 2010) (Figure 5A). To investigate the effect of elesclomol on mitochondrial Fe-S cluster biosynthesis, we utilized *in vitro* Fe-S cluster assembly assays. Assembly was initiated by the addition of cysteine and titration of elesclomol (Figure 5B) or its analog (compound-1) (Figure S5A) at 5X and 10X stoichiometry (relative to ISCU and FDX1) into the reaction mix progressively inhibited *in vitro* Fe-S cluster assembly. In contrast, substitution of reduced FDX1 by its homolog, reduced FDX2, resulted in almost complete loss of inhibition by elesclomol (Figure S5B). In addition, elesclomol showed no inhibitory effect on cysteine desulfurase activity with DTT as the reductant (Figure S5C). These results are all consistent with a direct inhibition of FDX1 function by elesclomol. As further confirmation, we examined the effect of elesclomol on electron transfer from reduced FDX1 to the ISC core complex upon the addition of cysteine. In this assay, elesclomol induced progressive inhibition of electron transfer from reduced FDX1 to the NFS1 component of the ISC core complex (Figure 5C).

**Figure 5.**
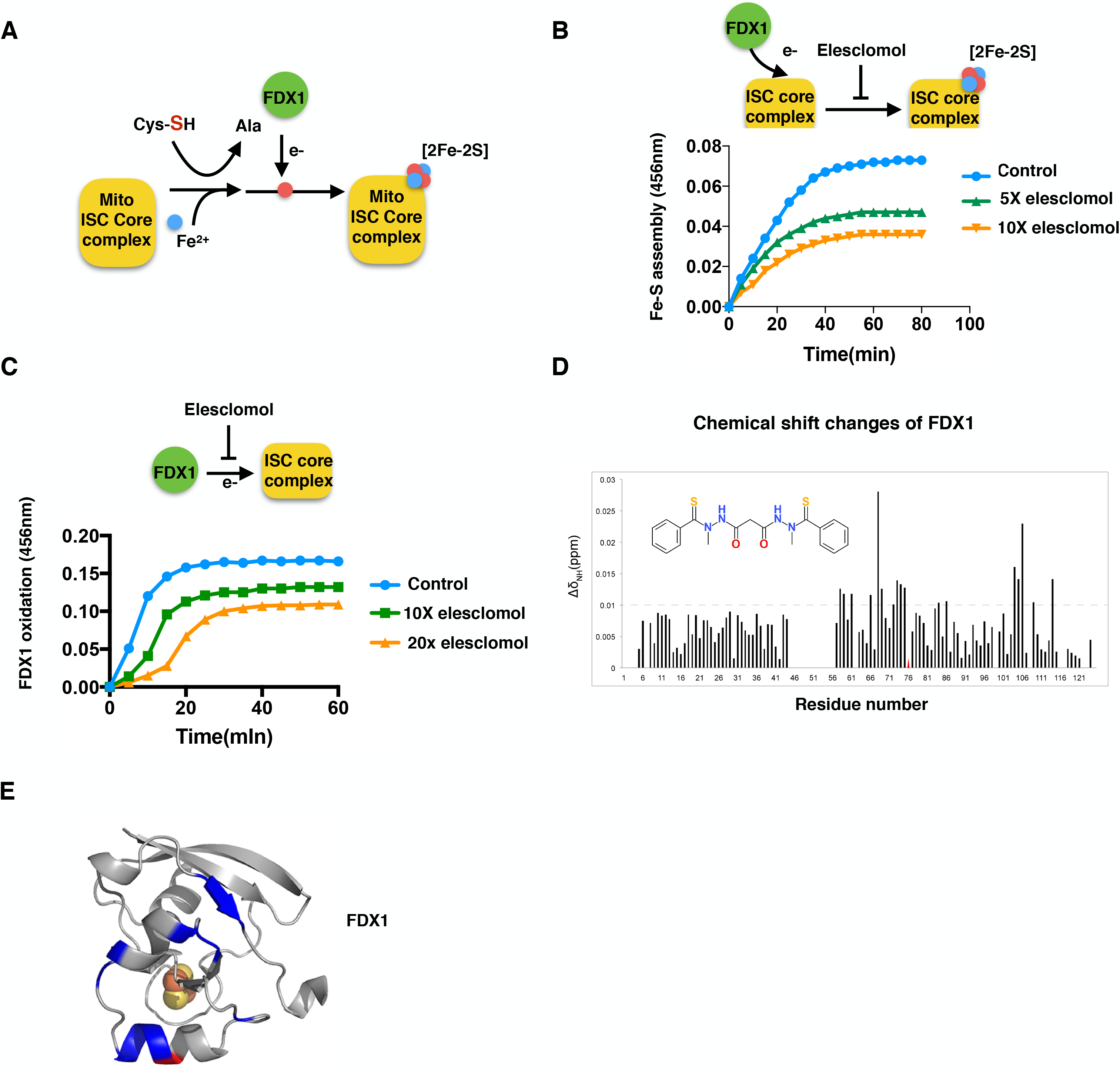
*In vitro* evidence that elesclomol directly inhibits mitochondrial iron sulfur (Fe-S) cluster biosynthesis by inhibiting electron transfer from FDX1 to cysteine desulfurase. (A) Schematic describing mitochondrial Fe-S cluster biosynthesis. The mitochondrial ISC (iron-sulfur cluster) core complex contains the scaffold protein ISCU and cysteine desulfurase (ACP-ISD11-NFS1)_2_. The latter catalyzes the conversion of cysteine to alanine and generates S^0^ for iron sulfur cluster assembly. S^0^ is reduced by FDX1. A [2Fe-2S] cluster is subsequently formed on ISCU. (B) Elesclomol inhibits *in vitro* Fe-S cluster assembly. *In vitro* Fe-S cluster assembly was carried out with reduced FDX1 as the reducing agent in the presence or absence of either 5X (green) or 10X (yellow) elesclomol (both relative to FDX1). Fe-S cluster formation was monitored by following the increase of absorbance at 456 nm. (C) Elesclomol inhibits electron transfer from reduced FDX1 to the cysteine desulfurase complex upon the addition of cysteine as demonstrated by the rate of oxidation of reduced FDX1. (D) Chemical shift (CS) perturbation (Δδ_NH_) analysis of [U-15N]-FDX1 upon interaction with elesclomol. The red triangle denotes an NMR peak that became severely broadened. (E) CS perturbation results from panel *D* mapped onto a diagram of the structure of FDX1. Color code: grey, not significantly affected (Δδ_NH_ < 0.01 ppm); blue, significant chemical shift changes (ΔδNH > 0.01 ppm); red, severe line broadening; grey, no assignments. The [2Fe-2S] cluster in FDX1 is indicated by spheres.

As further confirmation of this activity, we found that elesclomol directly binds FDX1. Using NMR spectroscopy, we showed that titration of elesclomol against [U-^15^N]-labeled FDX1 led to significant chemical shift changes in the 2D ^1^H,^15^N TROSY-HSQC spectrum (Figure S5D). Careful analysis of the chemical shift (CS) perturbations and peak broadenings revealed that the most affected FDX residues (Δδ > 0.01 ppm) corresponded to I58-F59, D61, K66, D68, A69, D72-D76, L84, and T104-R106 (Figure 5D). Most of these residues map to the α2, α3 helices and the β5 strand within the FDX1 structure (Figure 5E). Interestingly, these residues have been previously shown to interact with cysteine desulfurase (Cai et al., 2017b). Thus, elesclomol binds and directly inhibits FDX1 activity, possibly by disrupting the interaction between FDX1 and cysteine desulfurase. As a result, electron transfer from reduced FDX1 to the cysteine desulfurase complex is decreased, which ultimately inhibits Fe-S cluster assembly.

### FDX1 inhibition with elesclomol impedes resistance to proteasome inhibitors in a mouse model of multiple myeloma

Proteasome inhibitors such as bortezomib are used as front-line therapies for multiple myeloma. Unfortunately, patients often rapidly develop resistance to these agents. Our results indicate that proteasome inhibitors preferentially target cells in the glycolytic state. Therefore, these drugs may select for cells that either already reside in the OXPHOS-state or are able to rapidly shift into this drug-tolerant metabolic state. Elesclomol by targeting the mitochondrial iron sulfur pathway inhibits the ability of cells to grow in Hi-OXPHOS and prevents the Lo19S proteasome inhibitor resistance in culture (Figure 2 and 3). Therefore, we hypothesized that elesclomol treatment might increase the efficacy of proteasome inhibitors in a clinically relevant orthotopic mouse model of multiple myeloma cancer cells.

To test this hypothesis, we established an orthotopic luciferized MM.1.S xenograft model by intravenous injection of MM.1.S cells into severe combined immunodeficient (SCID) mice. After engraftment, the mice were randomized into treatment groups, each carrying equivalent tumor burdens based on bioluminescence imaging (BLI). Mice were then treated with either vehicle alone, bortezomib, elesclomol or with the two drugs in combination. A separate group was treated with bortezomib alone until tumor progression was documented by BLI (day 15) at which point elesclomol was added to the treatment regime (“delayed group”) (Figure 6A).

**Figure 6.**
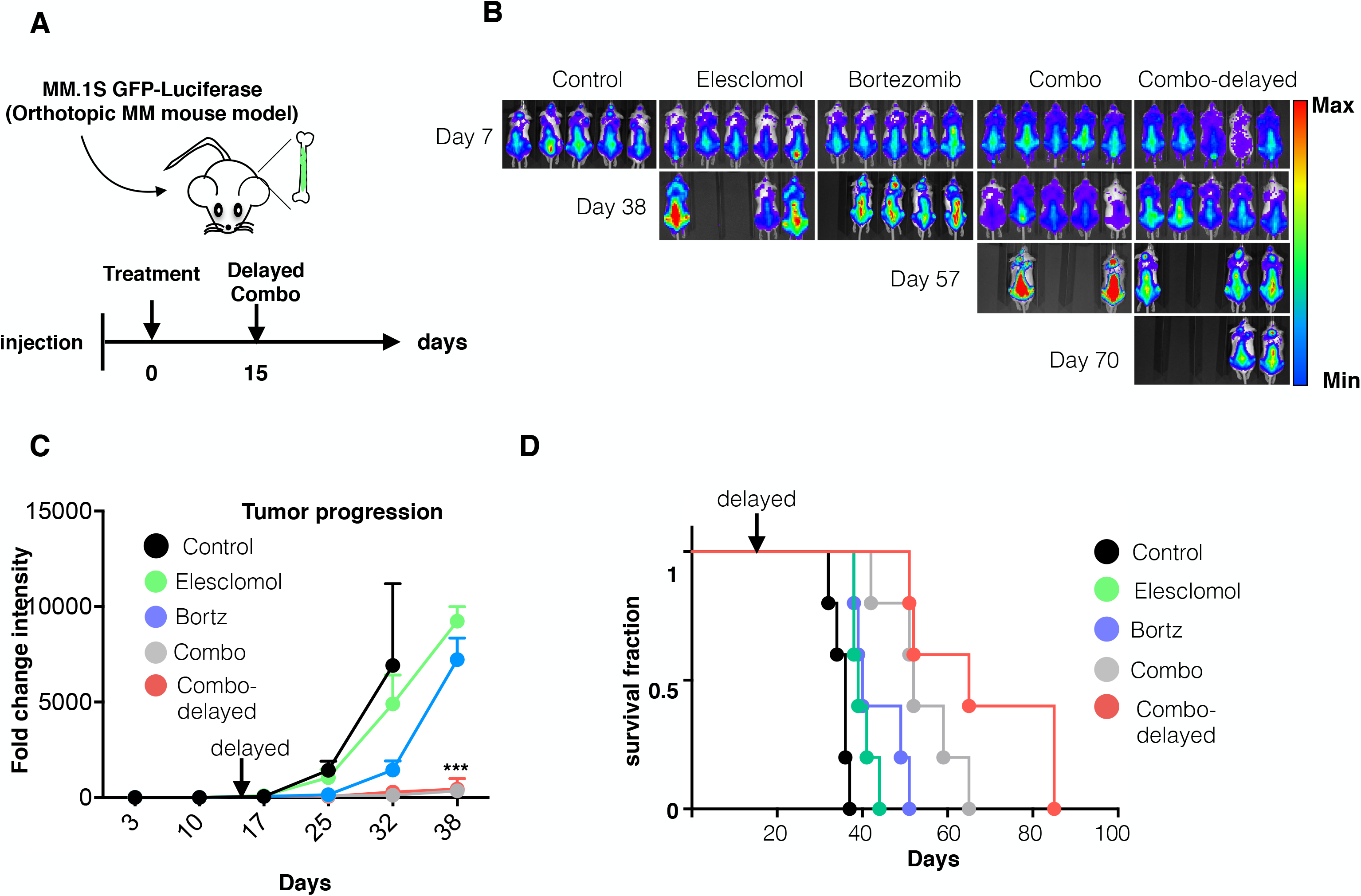
Inhibiting the glycolysis-OXPHOS shift with elesclomol is beneficial to proteasome inhibitor treatment in a mouse model of multiple myeloma. (A) Schematic of the experimental outline. Mice were injected with MM.1S luciferase expressing cells. Upon tumor formation as judged by BLI signal intensity, treatment with elesclomol (28mg/kg), bortezomib (0.25mg/kg) or the combination (Combo) was initiated (day 0). At day 15, elesclomol was added to the treatment of one of the groups that had received only bortezomib from day 0 to day 15 (combo-delayed group). (B) Representative *in vivo* images of MM1S LUC/GFP tumor-bearing SCID mice over the course of the indicated treatments. (C) Tumor burden over time as determined by changes from the baseline radiant flux associated with the BLI signal intensity (statistical test: Wilcoxon) (D) Kaplan Meier survival curve of MM1S tumor-bearing SCID mice; the black arrows indicate the timing of the elesclomol inclusion for the bortezomib+elesclomol delayed group (statistical test: Log-rank). N = 5 per group. *** p<0.001.

While bortezomib or elesclomol alone each had modest effects on tumor burden, BLI monitoring demonstrated marked inhibition of tumor progression in the elesclomol-bortezomib combination groups (Figure 6B-D), despite the very aggressive nature of this particular MM cell line. Even more importantly, we observed significant effects of the combination treatment on survival. Elesclomol treatment alone modestly but significantly increased survival (median OS 39 versus 36 p=0.0021). However, the strongest effects on survival were observed in the bortezomib-elesclomol combination groups: the combination treatment significantly increased survival relative to the control (median OS 52 versus 36 p=0.0021) and to bortezomib alone (median OS 52 versus 40 p=0.024). Interestingly, we observed the most striking treatment effect in the delayed bortezomib-elesclomol group, which exhibited the greatest improvement in survival over treatment compared with control (median OS 65 versus 36 p=0.0021) or bortezomib treatment alone (median OS 65 versus 40 p=0.0044). Thus, inhibiting FDX1 with elesclomol impaired the ability of multiple myeloma cells to cope with proteasome inhibitor-induced proteotoxic stress and markedly improved disease control in our orthotopic mouse model.

## DISCUSSION

The mechanisms responsible for acquired proteasome inhibitor resistance in hematological cancers and for the poor clinical activity of proteasome inhibitors against most solid tumors remain poorly understood. These challenges block the realization of the full clinical potential of proteasome inhibitors. Our work herein suggests that a cellular metabolic shift from glycolysis to OXPHOS is an integral part of a mechanism that cancer cells deploy to cope with the proteotoxic stress induced by proteasome inhibitors. Indeed, a frequent mechanism by which cancers acquire resistance to proteasome inhibitors, namely the Lo19S state, shows tight association with increased dependence on OXPHOS, which is readily detected as broad up-regulation of mitochondrial gene expression in many tumor types. Recapitulation of the naturally occurring Lo19S inhibitor resistant state through experimental 19S subunit knockdown markedly increases the ability of cancer cells in cell culture to withstand proteotoxic stress. However, inhibition of the mitochondrial iron-sulfur pathway using the first-in-class FDX1-specific inhibitor characterized herein, attenuates the cancer cell ability to cope with proteasome inhibitor-induced toxicity. This occurs not just in cell culture but more importantly, in a highly aggressive orthotopic mouse model of multiple myeloma.

The concept that shifts in core metabolism play an important role in anticancer drug-resistance has recently found support in multiple models (Ippolito et al., 2016; Kuntz et al., 2017; Lee et al., 2017; Matassa et al., 2016; Vazquez et al., 2013; Vellinga et al., 2015). However, the mechanism(s) by which shifts in metabolism are associated with numerous drug-resistant cancers are just beginning to be deciphered. One possibility is that mitochondrial functions are crucial to developing and sustaining the drug-resistant state. These functions extend beyond mitochondrial ATP production and include as metabolite production and redox homeostasis.

For example, mitochondrial Fe-S cluster biosynthesis is a process that is highly conserved in eukaryotic organisms from yeast to man (Lill and Muhlenhoff, 2006). In cancer cells, it has been reported that this biosynthetic pathway is essential for proliferation in the OXPHOS state (Arroyo et al., 2016) and for primary lung or metastatic tumors growing in high oxygen levels (Alvarez et al., 2017). This could be largely due to the numerous mitochondrial proteins that depend on Fe-S clusters for their functionality including complex I-III proteins, ACO1 and others making it a key upstream regulator of mitochondrial function (Cameron et al., 2011). However, the importance of the mitochondrial Fe-S cluster pathway extends beyond sustaining the electron transport chain and might also play a crucial role in regulating the cellular ROS-mediated cell death mechanisms such as ferroptosis (Alvarez et al., 2017). Indeed, conclusions from our genetic screens and *in vitro* characterization suggest that targeting mitochondrial Fe-S cluster biosynthesis may also provide a feasible strategy for overcoming proteasome inhibitor resistance.

Long before any of the mechanistic insights reported here, elesclomol underwent clinical development for the treatment of advanced solid tumors in combination with paclitaxel (O’Day et al., 2009; O’Day et al., 2013). Although the mechanism of action for elesclomol was not known it was suggested that elesclomol induced cytotoxicity is mediated by increased reactive oxygen specifies (ROS) levels (Kirshner et al., 2008) that might be a result of inhibition of the mitochondrial respiratory chain function (Barbi de Moura et al., 2012; Blackman et al., 2012). Here we show that elesclomol inhibits the Fe-S sulfur pathway acting as an upstream regulator of mitochondrial function. Our genetic screen also indicated that elesclomol-mediated toxicity is highly dependent on the expression of *FDX1*, a critical component in the Fe-S cluster synthesis pathway. Interestingly, *FDX1*’s close family member *FDX2* or other mitochondrial genes also essential for cancer cell growth in the OXPHOS state were not identified in our genetic screen. This fact and the remarkably strong and restricted selection for *FDX1* knockout cells in our CRISPR/Cas9 screen suggest a toxic gain of function exerted by FDX1 in the presence of elesclomol. Moreover, although both FDX1 and FDX2 are active as reductants in iron-sulfur cluster assembly (Cai et al., 2017b), our finding further suggests that there must be some critical electron transfer function fulfilled specifically by FDX1.

The Lo19S state is achieved by reduced gene expression of any one 19S subunit. However, other post-translational events can regulate the levels of intact 26S proteasomes including oxidative stress (Wang et al., 2010), NADH/NAD^+^ balance (Cho-Park and Steller, 2013; Tsvetkov et al., 2014), post-transcriptional regulation (Lokireddy et al., 2015; Myeku et al., 2016) and chaperone-mediated assembly (Kaneko et al., 2009; Rousseau and Bertolotti, 2016). Many of these processes are also associated with the cell metabolic state, which suggests that the dynamic interaction between metabolism and protein breakdown mediated by the proteasome could extend beyond transcriptional regulation of 19S subunits. In yeast, the shift from proliferative, glucose-dependent growth to a stationary respiring state is associated with a substantial decrease in the abundance of 26S proteasome complexes but not the 20S complex (Lo19S state) (Glickman et al., 1998). Thus, the dynamic interplay between regulation of the levels of the cellular proteasome complex regulation and metabolism can extend beyond cancer drug resistance to other models of normal development and aging.

Cancer cells often co-opt and re-wire adaptive non-oncogenic systems for their benefit. Here, we show that the Lo19S state is associated with Hi-OXPHOS and enables cells to withstand greater proteotoxic stress. The tight link between protein turnover and cellular metabolism is ancient and evolutionarily conserved, deriving from a requirement of cells to coordinate energy production with protein homeostasis. This feedback loop stems, on the one hand, from the energy requirements of protein synthesis (Frumkin et al., 2017) and breakdown (Peth et al., 2013) and, on the other hand, the need to recycle damaged, oxidized, dysfunctional proteins and enzymes (Raynes et al., 2016). The proteasome plays a key role in linking these processes by controlling the recycling of amino acids, which are the building blocks of protein synthesis and key intermediaries in many metabolic and redox pathways (Suraweera et al., 2012; Vabulas and Hartl, 2005). Thus, from a broader evolutionary perspective, combined targeting of two evolutionary conserved pathways, the protein degradation pathway and the mitochondrial Fe-S cluster pathway is an attractive therapeutic strategy as it confronts the cancer cells with opposing selective pressures. Elevated proteasome function is required for the glycolytic proliferating cancer cells and can be targeted with proteasome inhibitors. Increased mitochondrial function is favorable for proteasome inhibitor drug-resistance but entails major sensitization to targeted inhibition of the mitochondrial Fe-S cluster synthesis pathway with elesclomol. As supported by our findings in mice, creating such a dilemma offers intriguing promise as a resistance-evasive anticancer strategy.

## Acknowledgments

This work is dedicated to the memory of Susan Lindquist who served as a great inspiration as a scientist,, mentor and a human being. We thank Linda Clayton, Can Kayatekin, Brooke Bevis, Naama Kanarek (Whitehead Institute) and Neekesh Dharia (Broad institute) for constructive discussion and comments. Special thanks to David Sabatini (Whitehead Institute) for overseeing this work and for providing critical comments and reviewing of the manuscript and Gerry Fink (Whitehead Institute) for his supervision. We thank the Koch Institute Swanson Biotechnology Center for technical support, specifically Jaime Cheah for her support conducting the chemical drug screen. NMR spectroscopy was carried out at the National Magnetic Resonance Facility at Madison which is supported by National Institutes of Health (NIH) grant P41GM103399; other work at the University of Wisconsin-Madison was supported by funds from the Biochemistry Department. P. Tsvetkov was supported by EMBO Fellowship ALTF 739-2011 and by the Charles A. King Trust Postdoctoral Fellowship Program. S.L. was an investigator of the Howard Hughes Medical Institute.

## Author Contributions

Conceptualization, P.T.; Investigation P.T., A.D., K.C., H.R.K,; Formal analysis, P.T., Pr.T., G.K., H.R.K.; Resources-Z.B. W.Y., A.T.; Writing original-draft P.T. Writing review & editing-P.T., H.R.K, S.S., L.W., J.L.M.; Funding acquisition S.L., J.L.M, I.M.G.; Supervision S.L, L.W., J.L.M, I.M.G;

## Materials and methods

### TCGA analysis

TCGA data analysis-The Cancer Genome Atlas (TCGA: <u>cancergenome.nih.gov</u>) expression (RNASeq V2) were downloaded using TCGA-assembler (Zhu et al., 2014). RNASeq data were quantified as RSEM. Sigma score was calculated for each primary tumor category separately by calculating a Z-score for every individual proteasome subunit gene and categorizing the tumors as 3-sigma or control as previously described (Tsvetkov et al., 2017).

Enrichment analysis was performed using GSEA (Subramanian et al., 2005), using H and C2 genesets of MSigDB. GO enrichment was visualized using ClueGO (Bindea et al., 2009) via Cytoscape (Shannon et al., 2003).

### Elesclomol sensitivity and gene dependency analysis

Dose-response and area under the curve metrics for 632 cell lines in response to treatment with elesclomol were obtained from the Genomics of Drug Sensitivity in Cancer (http://www.cancerrxgene.org/downloads) (Yang et al., 2013). Results of genome-scale CRISPR/Cas9 knockout viability screens for 17,673 genes in 391 cell lines were obtained from the Cancer Dependency Map (https://depmap.org). The relationship between CRISPR knockouts, response to elesclomol was assessed using the limma R package (Ritchie et al., 2015). The relationship between elesclomol response from GDSC and the CRISPR knockouts was assessed across 246 common lines in both datasets while the relationship between FDX1 knockout and CRISPR knockouts was assessed across all 391 lines present in the knockout dataset.

### Antibodies reagents

FDX1 (Proteintech Group, Inc (Catalog Number: 12592-1-AP)), tubulin (ab80779 Abcam)

### Compounds

elesclomol (MedChemexpress Co., LTD. # HY-12040), Ixazomib, disulfiram and antimycin A (selleck), Bortezomib (LC-Laboratories). Elesclomol analogs (OnTarget Pharmaceutical Consulting LLC.) Drug libraries used were the anti cancer compound library L3000 (selleck), BML-2865 Natural product library (Enzo), the NIH Clinical Collections (NCC) and the Boston University’s Chemical Methodology and Library Development (CMLD-BU) drug library (http://www.bu.edu/cmd/about-the-bu-cmd/compound-libraries/).

### Cell culture methods

T47D, Lo19S T47D, K562 and 293T-HS cells were cultured in RPMI-1640 medium supplemented with 10% fetal bovine serum; HEK293, HEK293T and MCF-7 cells were cultured in Dulbecco’s modified Eagle’s medium supplemented with 10% fetal bovine serum. For the glucose, galactose experiments media lacking glucose was supplemented with dialyzed serum and either 10mM Glucose or 10mM galactose.

GFP^+^/Luc^+^ MM.1S cells were generated by retroviral transduction of the human MM1.S that was purchased from ATCC (Manassas, VA, USA)., using the pGC-GFP/Luc vector. Cells were authenticated by short tandem repeat DNA profiling.

Generation of the Lo19S T47D cell line – For the generation of the T47D Tet-inducible PSMD2 knockdown cell line-the TRIPZ vector with an inducible shRNA targeting PSMD2 was purchased from Dharmacon (clone V3THS_403760). It was introduced to the T47D cells according to manufactures protocol and cells were selected with puromycin 1mg/ml for one week. The cells were exposed to doxycycline for 24 hours and cells were FACS sorted for the top 10% of most RFP expressing cells (highest expression of shRNA). The cells were further cultured in the absence of doxycycline and PSMD2 knockdown was induced as specified in the text.

### The Lo19S T47D drug screen

Cells with a Dox inducible PSMD2 KD plasmid were incubated in the presence or absence of 1ug/ml of Doxycycline for 48 hours (control versus PSMD2 KD respectively). After 48 hours the cells were collected counted and plated (in the absence of dox) at 1000 cells/well in 384-well opaque, white assay plates (Corning, NY), 50 uL per well, and incubated overnight at 37°C/5% CO2. Compound stocks from the Cancer drug library containing 349 bioactive compounds that were arrayed in dose 10nM, 100nM, 1uM, 10uM (Selleck anti-cancer compound library L3000), the natural products library containing 502 compounds that were utilized in 5 doses (dilution of 1:1000 of 2ug/ml, 0.2ug/ml, 0.002/ug/ml, 0.0002ug/ml), the NIH bioactive library including 731 drugs in dose of 10nM, 100nM, 1uM, 10uM and the Boston University’s CMLD (Chemical Methodology and Library Development) compound deck containing 2866 compounds in one dose (10uM), including novel chemotypes that uniquely probe three-dimensional space by employing stereochemical and positional variation within the molecular framework as diversity elements in library design. 50 nl of compounds were pin-transferred (V&P Scientific, CA, pin tool mounted onto Tecan Freedom Evo 150 MCA96 head, Tecan, CA) into duplicate assay plates and incubated for 72h. The DMSO content was 0.1% within each well. Per plate, there are 32 wells of DMSO vehicle control and 32 wells of positive control compounds. After three days of incubation, 10 uL of CellTiter-Glo (Promega, WI) was added to each well, incubated for 10 minutes and the luminescence output was read on the M1000 Infinite Pro plate reader (Tecan, CA). CellTiter-Glo measures ATP levels in the cell and is used as a surrogate for cell viability.

### The glucose/galactose drug screen

T47D cells growing in regular media were collected counted and after centrifugation cells were re-suspended in RPMI media without glucose containing 10% dialyzed serum with the addition of either 10mM glucose or 10mM galactose. Cells were then plated at 1000 cells/well in 384-well opaque, white assay plates (Corning, NY), 50 uL per well, and incubated overnight at 37°C/5% CO2. Application of the drug libraries and cell viability was executed as described above for the Lo19S drug screen

### Heat-shock reporter

293T cells (American Type Culture Collection) harboring a enhanced GFP fused to firefly luciferase under control of HSP70B′ promoter elements as previously described (Wijeratne et al., 2014)

### CRISPR screen

268M cells were transduced with 22mL human genome-wide cleavage-optimized lentiviral sgRNA library containing Cas9 (http://www.addgene.org/pooled-library/sabatini-crispr-human-high-activity-3-sublibraries) as previously described (Wang et al., 2016). Cells were allowed to recover for 48h and selected with 3ug/mL puromycin for 72h and transduction efficiency was determined. 50M cells per condition were passaged every 2-3 days throughout the duration of the screen (except where noted), according to the following timeline: STA5781:

Day 0-7: 4nM

Day 7-26: 6nM

Day 26-34: 100nM

STA3998:

Day 0-7: 25nM

Day 7-23: 44nM

Day 26-31: 1uM

80M cells were collected after puromycin selection, representing the initial cell population, and 5-80M cells were collected at each time point. Genomic DNA was isolated using the QIAmp DNA Blood Maxi, Midi, or Miniprep kit, depending on the cell number of the sample, and high-throughput sequencing libraries were prepared REF (https://www.ncbi.nlm.nih.gov/pubmed/26933250), except that the following forward PCR primer was used:

AATGATACGGCGACCACCGAGATCTACACGAATACTGCCATTTGTCTCAAGATCTA

Sequencing reads were aligned to the sgRNA library and the abundance of each sgRNA was calculated. The counts from each sample were normalized for sequencing depth after adding a pseudocount of one. sgRNAs with fewer than 50 reads in the initial reference dataset were omitted from downstream analyses. The log2 fold change in abundance of each sgRNA between the final and initial reference populations was calculated and used to define a CRISPR Score (CS) for each gene. The CS is the average log2 fold change in abundance of all sgRNAs targeting a given gene. Genes represented by fewer than 5 sgRNAs in the initial reference dataset were omitted from downstream analyses.

### Statistical analysis

The distribution of the log2 fold changes between the final and initial populations for the set of sgRNAs targeting each gene was tested against the distribution for all sgRNAs by the Kolmogorov-Smirnov test. P-values were adjusted for multiple comparisons by the Benjamini-Hochberg (FDR) procedure.

### FDX1 gene targeting using single sgRNAs

plentiCRISPRv2 vector (Wang et al., 2016) was used as a backbone to insert the gRNAs targeting FDX1 and AAVS1 as control.

BsmBI restriction enzyme digest of the backbone vector was followed by T4 ligation of the digested vector with phosphorylated an annealed oligo pairs:

sgFDX1-1 **CACCG**GCAGGCCGCTGGATCCAGCG

sgFDX1-1_RC: **AAAC**CGCTGGATCCAGCGGCCTGC**C**

sgFDX1-2 **CACCG**TGATTCTCTGCTAGATGTTG

sgFDX1-2_RC: **AAAC**CAACATCTAGCAGAGAATCA**C**

sgAAVS1: **CACCG**GGGGCCACTAGGGACAGGAT

sgAAVS1_RC: **AAAC**ATCCTGTCCCTAGTGGCCCC**C**

To generate the lentiviruses, 3 × 106 HEK-293T cells were seeded in a 10 cm plate in DMEM supplemented with 10% IFS. 24 hours later, the cells were transfected with the above sgRNA pLenti-encoding plasmids alongside the Δ VPR envelope and CMV VSV-G packaging plasmids using the XTremeGene 9 Transfection Reagent (Roche). 12 hours after transfection, the medium was aspirated and replaced with 8 ml fresh medium. Virus-containing supernatants were collected 48 hours after the transfection and passed through a 0.45 µm filter to eliminate cells. Transduction of cells was done as described above.

K562 cell were cells were seeded at a density of 300,000 cells/mL in 6-well plates in 2 mL of RPMI containing 8 µg/mL polybrene (EMD Millipore), and then transduced with lentivirus by centrifugation at 2,200 RPM for 90 min at 37° C. After a 24-hour incubation, cells were pelleted to remove virus and then re-seeded into fresh culture medium containing puromycin, and selected for 72 hours. Cells were then single cell FACS sorted into 96 well plates and grown out in the presence of 1ug/ml puromycin.

### Viability and cell proliferation assays

Relative cell number count protocol-The viability of K562 cells was conducted by seeding cells at 250,000 cells/mL in 6-well plates of RPMI. In the presence or absence of different concentrations of compound-1 or −2. Viable cell numbers were counted using countess (Invitrogen) every 2-3 days and cells were reseeded at 250,000 cells/mL in 6-well plates of RPMI to continue growth.

The viability of FDX1 KO K562 cells and AAVS1 KO and WT controls was examined by plating 100,000 cells/mL in 6-well cells in RPMI media (without glucose containing dialyzed serum) with either 10mM glucose or 10mM galactose. Viable cell numbers were counted every day using countess (Invitrogen).

Cell viability-where indicated relative cell number following different drug treatment was conducted by plating cells at 1000 cells/well in 384 well plates. Indicated concentrations of compounds were added 24 hour after plating at least in triplicate for each condition. viability was measured 72 hours after drug addition using cell titer-glo (Promega) according to manufactures protocol. Same protocol was applied for the experiments conducted for the Lo19S T47D, MCF7 and T47D in the context of altered carbon source only the cells were plated at 1000 cells/well in RPMI media (without glucose and with dialyzed serum) containing either 10mM Glucose or 10mM galactose.

### Protein Expression and Purification

Unlabeled samples of ISCU, unlabeled (NFS1-ISD11-Acp)_2_ (abbreviated as SDA), unlabeled and [U-^15^N]-FDX1 (truncated first 61 amino acids) and unlabeled FDX2 were produced and purified as described previously (Cai et al., 2013; Cai et al., 2017a; Cai et al., 2017b). FDX1 and FDX2 were reduced by adding a 10-fold excess of sodium dithionite to oxidized FDX1 or FDX2 in an anaerobic chamber (Coy Laboratory). The reduction of FDX1 or FDX2 was monitored by the color change from dark brown to light pink and by recording UV/vis spectra before and after reduction. The sample was dialyzed extensively against anaerobic HN buffer to remove excess sodium dithionite.

### NMR Spectroscopy

NMR spectra were collected at the National Magnetic Resonance Facility at Madison on a 750 MHz (^1^H) Bruker with a *z*-gradient cryogenic probe. The buffer used for NMR samples (HNT buffer) contained 20 mM HEPES at pH 7.6, 150 mM NaCl, and 2 mM TCEP. All sample temperatures were regulated at 25 °C. NMRPipe software (Delaglio et al., 1995) was used to process the raw NMR data and NMRFAM-SPARKY (Lee et al., 2015) software was utilized to visualize and analyze the processed NMR data.

To study the interactions of FDX1 with elesclomol, 0.2 mM [U-^15^N]-FDX1 in HNT buffer were placed in 5 mm Shigemi NMR tubes, and 2D ^1^H,^15^N TROSY-HSQC spectra were collected before and after titration of 5 equivalent molar ratio of elesclomol dissolved in HNT buffer.

Chemical shift perturbations (Δδ_HN_ absolute value ppm) were calculated by

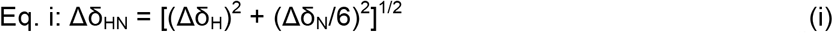

where Δδ_H_ and Δδ_N_ are the chemical shift changes in the ^1^H and ^15^N dimensions, respectively.

### Cysteine desulfurase assay

The protein samples used in the cysteine desulfurase assay and Fe-S cluster assembly experiments were prepared in an anaerobic chamber (Coy Laboratory) with samples buffer-exchanged extensively prior to the experiments with anaerobic buffer containing 20 mM HEPES at pH 7.6 and 150 mM NaCl (HN buffer). The reaction volumes in all the experiments were kept to 1 ml. A Shimadzu UV-1700 uv/visible spectrophotometer with a temperature control unit was used to collect the spectra, and UVProbe 2.21 software (Shimadzu) was used in collecting and analyzing the data.

The cysteine desulfurase assay reactions (300 µL in HN buffer) contained 10 µM SDA. The reductant was 100 µM DTT. 100 µM L-cysteine was added to initiate the reaction. 10 to 200 µM elesclomol (1-20 X relative to SDA) was added to assess their effect on sulfide production. After 20 min of anaerobic incubation at room temperature, the reaction mixture was diluted to 800 µL, and 100 µL of 20 mM N,N-dimethyl-p-phenylenediamine in 7.2 M HCl and 100 µL of 30 mM FeCl_3_ in 1.2 M HCl were added to quench the reaction and convert sulfide to methylene blue. The quenched reaction was incubated for 15 min at room temperature, and then the absorbance at 670 nm was measured and used to estimate the amount of sulfide by comparison to a standard curve obtained from known concentrations of Na_2_S.

### In *vitro* Fe-S cluster assembly assay

The *in vitro* Fe-S cluster assembly assays were carried out as follows. Reaction mixtures (1 mL) prepared in the anaerobic chamber contained 25 µM reduced FDX1 or reduced FDX2 as the reductant, 1 µM SDA, 25 µM ISCU and 100 µM (NH_4_)_2_Fe(SO_4_)_2_. To test the effect of the drug, reaction mixtures contained 25-250 µM elesclomol (1-20 X relative to FDX1). L-cysteine (at a final concentration 100 µM) was added to initiate each experiment. Samples were then transferred to 1-cm path-length quartz cuvettes, sealed with rubber septa, and uv/vis spectra were collected at 25 °C. The growth of absorbance at 456 nm was used to monitor the assembly of [2Fe-2S] clusters.

### Electron transfer assay

Electron transfer from re-FDX1 SDA complex was monitored as follows. 25 µM reduced FDX1 was mixed with 25 µM SDA, and 125 µM L-cysteine was added to initiate the reaction. To test the effect of the drug, samples contained 25-500 µM elesclomol (1-20 X relative to FDX1). Samples were then transferred to 1 cm path-length quartz cuvettes, sealed with rubber septa, and UV/vis spectra were collected at 25 °C. The growth of absorbance at 456 nm was used to monitor the oxidation of reduced FDX1 as a result of electron transfer to SDA.

### Disseminated Multiple Myeloma Animal model and survival study

A total of 25 Female SCID-beige mice at 4-6 weeks of age (Taconic, USA) were utilized for the study. Tumors cells (3 million LUC^+^/RFP^+^ MM1S cells/mouse) were introduced into mice via intravenous injection and were allowed to grow until the LUC signal from their tumors reached 1×10^7^ radians (photons/sec/cm^2^/surface area). Tumor growth was monitored weekly by bioluminescence imaging (BLI), using an IVIS Spectrum-bioluminescent and fluorescent imaging system (Perkins Elmer). Mice were randomly attributed a group of treatment (n=5 per group). Control group was IV injected once a week 200 uL saline solution; Bortezomib (0.25 mg/kg, 100 uL) was IP injected first day of the week, once a week until death of the animal. Elesclomol (25 mg/kg, 100 uL) was subcutaneously injected every 3^rd^ day of the week until death of the animal. Elesclomol+Bortezomib group were treated similarly to previous groups. The delayed group started the Elesclomol treatment 14 days after the inclusion of the animal in the study. Endpoint of the study was dictated by either limb-paralysis of the animal or severe toxicity induced by the treatment as *per* Dana-Farber Animal policy.

---

**Figure S1.**
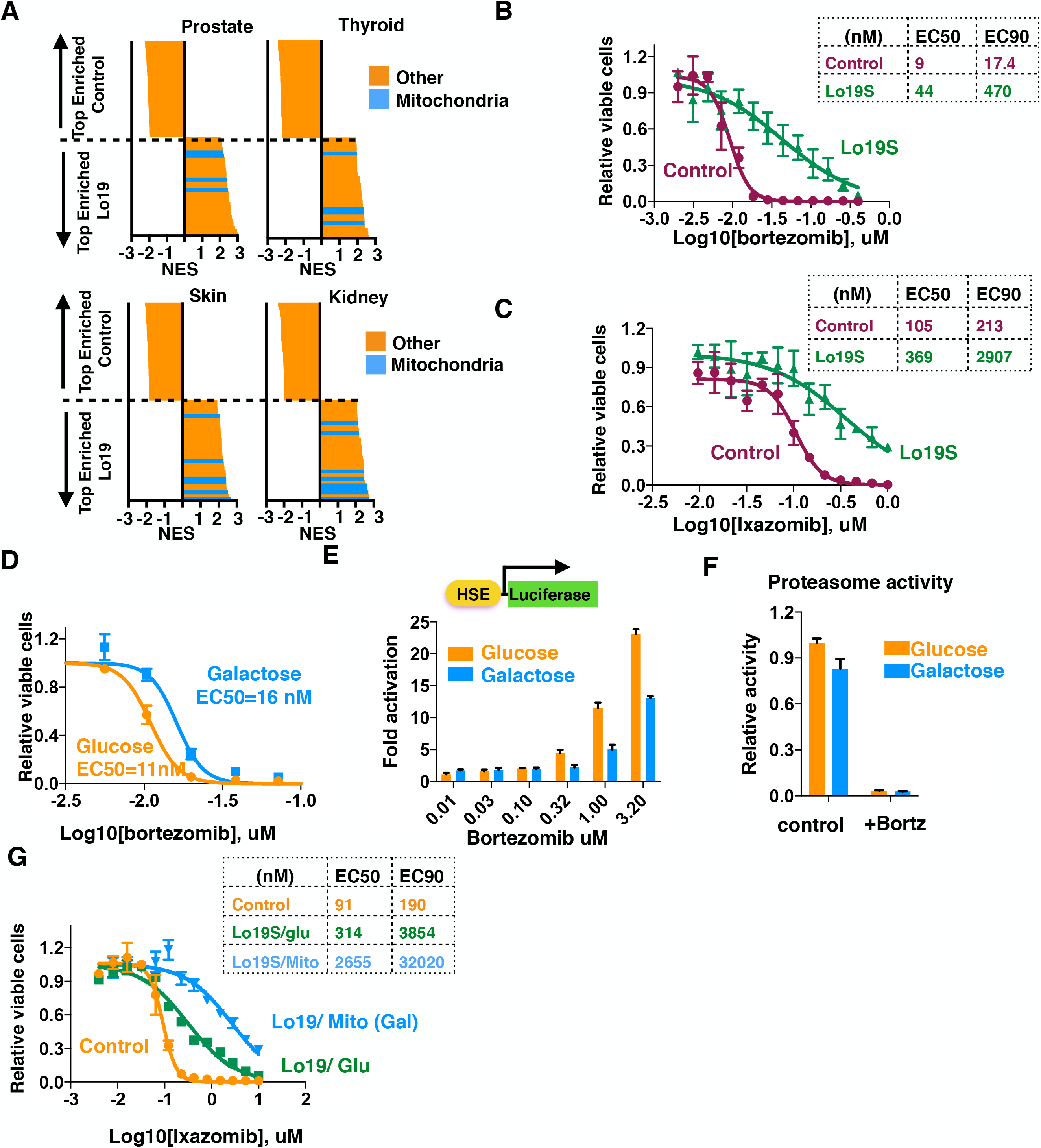
(A) Gene expression in cancers in the TCGA dataset was analyzed and stratified by tumors in the Lo19S state (one subunit of the 19S proteasome complex suppressed by more than 3 standard deviation (ref)) or control (the rest of the tumors). Gene set enrichment analysis (GSEA) of genes upregulated in Lo19S but not control tumors was conducted for prostate, thyroid, skin and kidney cancers from the TCGA. The top and bottom 29 categories are plotted. Mitochondrial-associated categories are marked in blue, the rest in orange. (B-C) The effect of induced Lo19S state on proteasome inhibitor resistance. T47D breast cancer cells harboring a doxycycline-inducible PSMD2 shRNA were grown in the presence or absence of 0.2 μg/ml doxycycline for 72 hours to induce the Lo19S state. Cells were then collected, washed and plated in the absence of doxycycline, 24 hour later either bortezomib (B) or ixazomib (C) were added at indicted concentrations and the relative cell number was measured 72 hours later. The calculated EC50 and EC90 are also plotted. (D) parental T47D breast cancer cells were examined for their relative viability when grown in the presence of bortezomib and media containing either glucose (control) or galactose (Hi-OXPHOS). (E) 293T cells harboring a heat-shock element (HSE) promoter followed by luciferase were examined for the ability of increasing concentrations of bortezomib to induce heat shock when cells were grown in media containing either glucose (control) or galactose (Hi-OXPHOS) (F) The chymotrypsin-like activity of the proteasome was determined in T47D cells grown in the presence of either glucose (control) or galactose (Hi-OXPHOS) with or without pre-treatment for 6 hours with 100nM bortezomib (+Bortz). (G) T47D breast cancer cells harboring a doxycycline-inducible PSMD2 shRNA were grown in the presence or absence of 0.2 μg/ml doxycycline for 72 hours to induce the Lo19 state. Cells were then collected, washed and plated in the absence of doxycycline in media containing either glucose (Glu) or galactose, which induces mitochondrial respiration (Mito). The relative viability was measured 72 hours after addition of the indicated concentrations of ixazomib.

**Figure S2.**
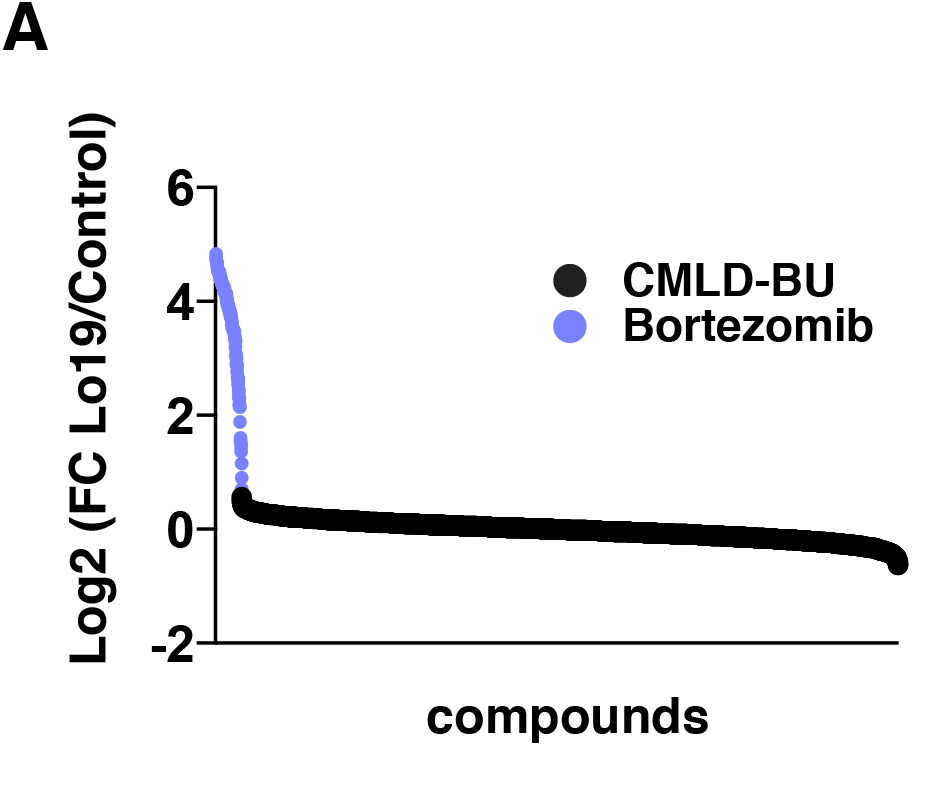
(A) T47D inducible Lo19S state cells and control cells (as described above) were subjected to 2866 compounds from the Boston University’s Chemical Methodology and Library Development (CMLD-BU) in one dose (10uM) and bortezomib as a control. The relative viability of the Lo19S cells versus control cells was calculated for replica experiments and the log2 of the ratio is plotted. Black dots: drugs in the library; purple dots: bortezomib controls.

**Figure S3.**
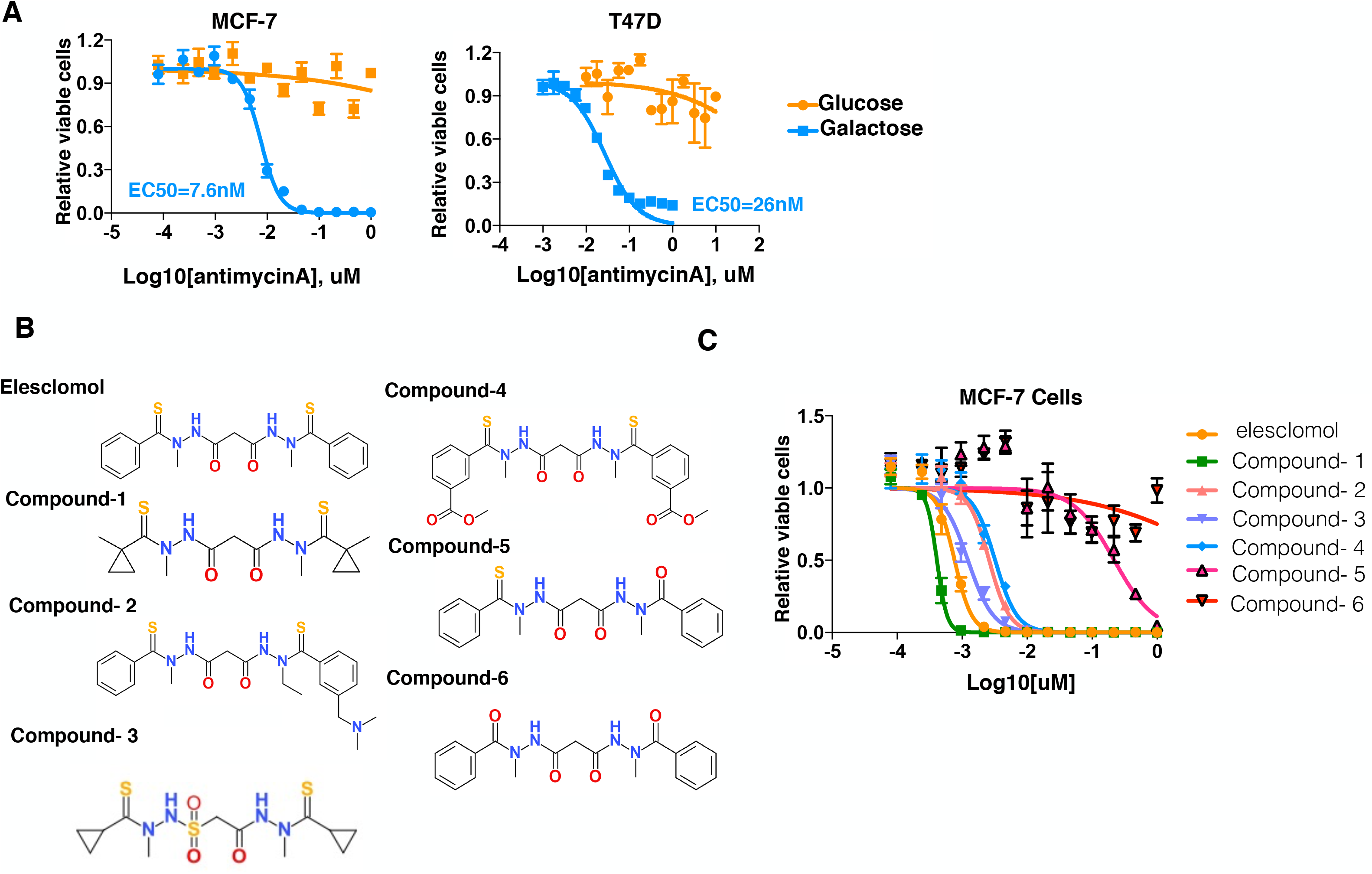
(A) MCF7 and T47D cells were grown in the presence of either glucose (control) or galactose (hi-OXPHOS) as the carbon source and the relative cell number was analyzed 72 hours after the addition of indicated concentrations of antimycin A. (B) the structures of the different elesclomol analogs used. Analogs named compound 1-6. (C) MCF-7 cells were grown in the presence of galactose (Hi-OXPHOS) and exposed to different elesclomol analogs (compounds 1-6) and relative cell growth (color coded) was analyzed 72 hours post compound addition. Plotted are the mean +-SD of at least three replicas and the calculated EC50s (A and C)

**Figure S4.**
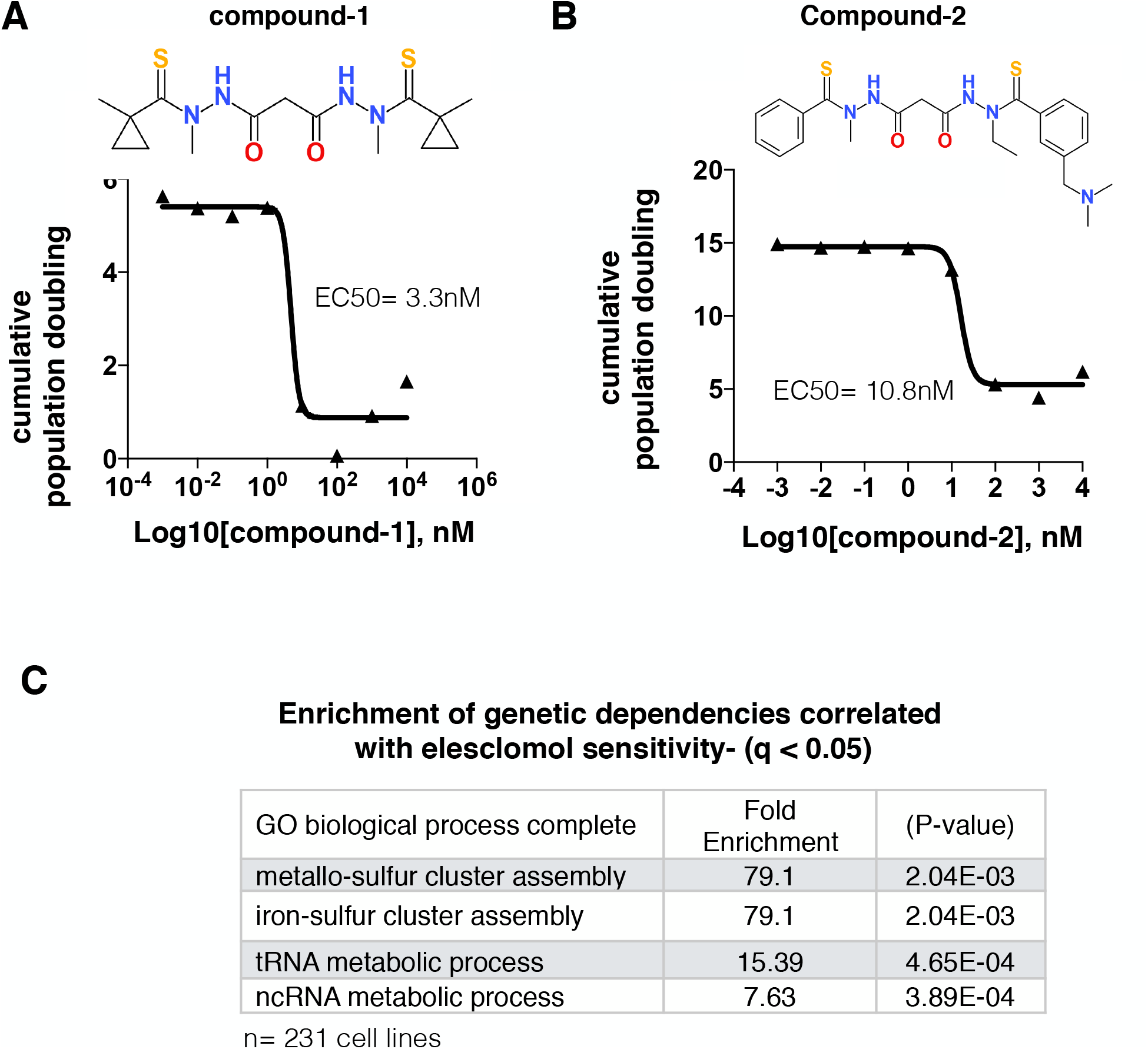
(A-B) The effect of the indicated concentrations of compound-1 (A) or compound-2 (B) on accumulative cell replications over 8 days. (C) Examining the CCLE dataset for correlation between the dependency of a gene (CRISPR score from CCLE) and sensitivity to elesclomol (taken from the GDSC). Analysis was conducted on 246 common cell lines in both datasets. GO ontology analysis was conducted on all genes that had a q value> 0.05 and the top hits are presented.

**Figure S5.**
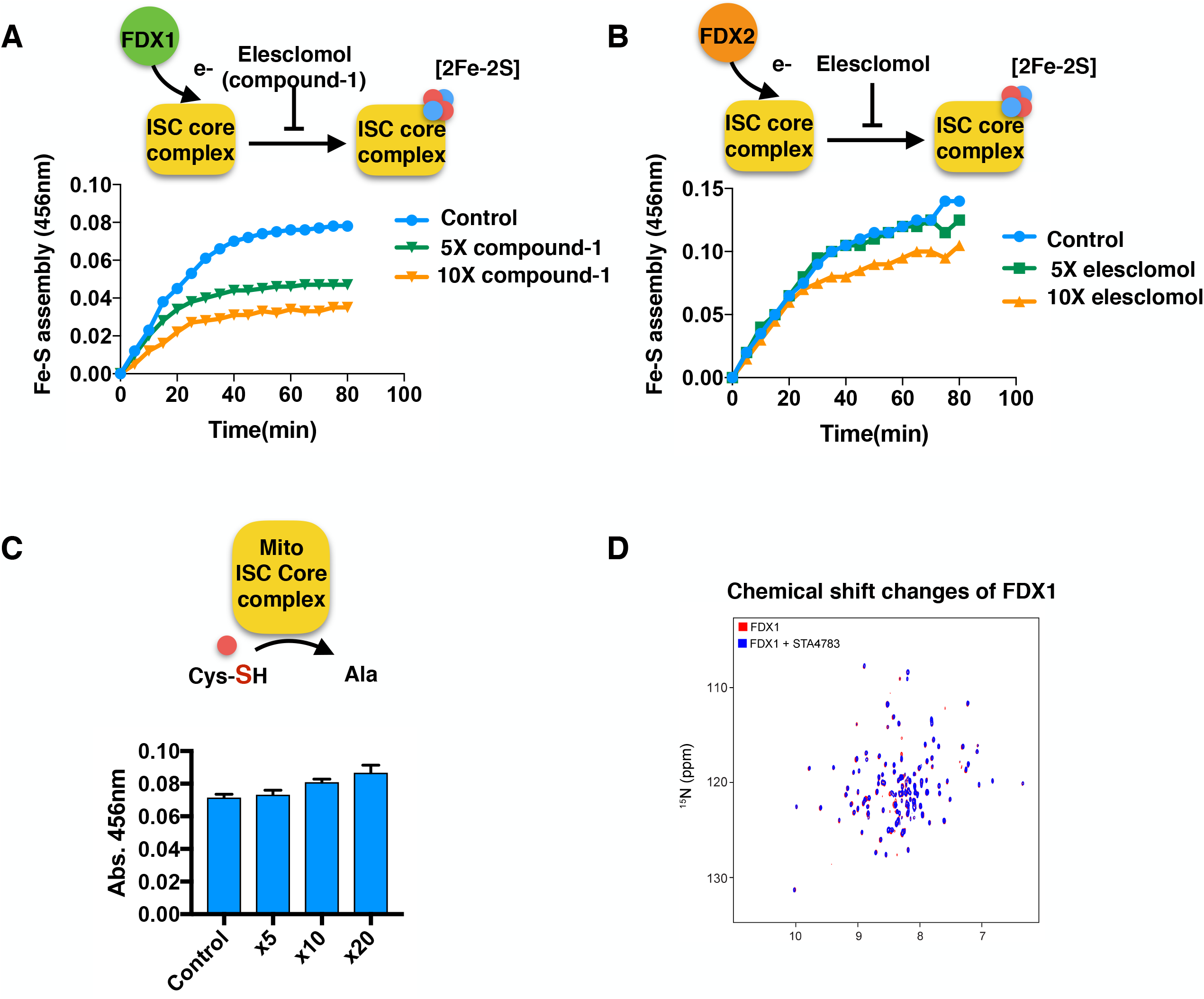
(A) Compound-1 (an elesclomol analog) significantly inhibits *in vitro* Fe-S cluster assembly with reduced FDX1 as the reducing agent. The *in vitro* Fe-S cluster assembly was carried out with reduced FDX1 as the reducing agent and detected by following the increase of absorbance at 456 nm following the addition of either of 5X (green) or 10X (yellow, both relative to FDX1) compound-1. (B) *In vitro* Fe-S cluster assembly was carried out with reduced FDX2 (instead of FDX1) as the reducing agent in the presence or absence of 5X (green, relative to FDX1) and 10X (yellow, relative to FDX1) elesclomol. Fe-S formation was monitored by following the increase of absorbance at 456 nm. (C) Elesclomol does not inhibit the activity of cysteine desulfurase. Cysteine desulfurase activity assay carried out with the indicated concentrations of elesclomol. (D) ^1^H,^15^N TROSY-HSQC NMR spectra of [U-^15^N]-FDX1 before (red) and after (blue) titration of 5X unlabeled elesclomol.

**Figure S6.**
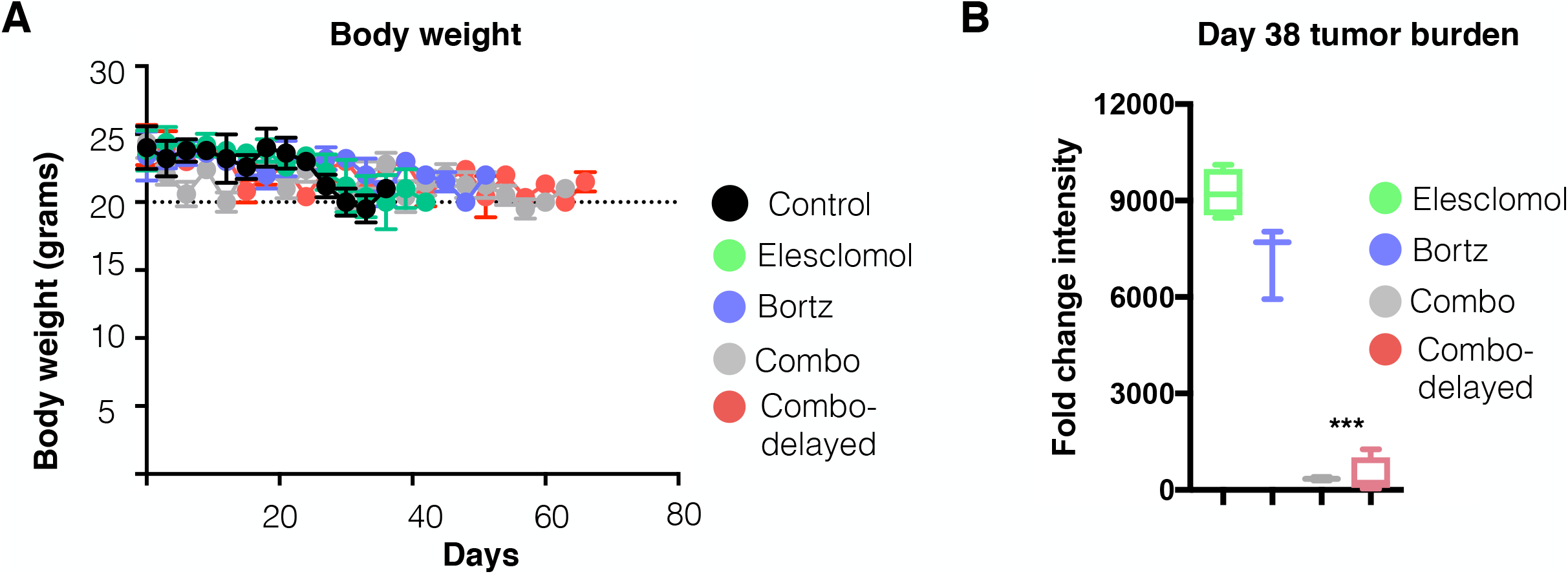
(A) Body weight of mice in the experimental set up described in figure 6. (B) Size of tumors in mice in the experiment described in figure 6 as assessed by BI measurement at week 38.

